# Interferon Restores Antigen Presentation and Sensitizes Medulloblastoma to T Cell Killing

**DOI:** 10.1101/2025.09.30.679624

**Authors:** Tanja Eisemann, Meher Beigi Masihi, Theophilos Tzaridis, Veronika Pister, Isaac Youm, Kendall R. Chambers, Aditi Dutta, Alexander T. Wenzel, Koei Chin, Scott L. Pomeroy, Jill P. Mesirov, Ernest Fraenkel, Anindya Bagchi, Lukas Chavez, Robert J. Wechsler-Reya

## Abstract

Medulloblastomas are commonly considered immunologically cold and refractory to immunotherapy. One contributing factor to their low immunogenicity is impaired antigen presentation, which allows tumor cells to escape from cytotoxic T cells. Here we use a syngeneic mouse model of medulloblastoma to study the role of CD8^+^ T cells in medulloblastoma growth. We demonstrate that despite low expression of MHC Class I on tumor cells, depletion of CD8^+^ T cells accelerates tumor growth, whereas adoptive transfer of tumor-reactive CD8^+^ T cells prolongs survival. These anti-tumor effects rely on T cells secreting interferon gamma (IFNγ), which induces MHC class I on tumor cells and facilitates tumor cell killing by T cells. Notably, this response is essential for CD8^+^ T cell–mediated tumor attack, as blocking IFNγ signaling in vivo abrogates MHC class I induction and eliminates the beneficial effect of T cells. Importantly, delivering IFNγ directly into tumors via convection-enhanced delivery (CED) enhances CD8^+^ T cell-mediated killing of tumor cells and significantly prolongs survival in tumor-bearing mice. These studies highlight the importance of T cells in controlling brain tumor growth and the value of IFNγ as an adjuvant for T cell-based immunotherapy.

## INTRODUCTION

Immunotherapy is a promising approach to treatment of cancer. Advances in our understanding of T cell biology, coupled with their innate physiological traits such as antigen-specificity and cytotoxicity, have made T cells a focal point in the development of immunotherapy, including immune checkpoint blockade, adoptive T cell therapy, and cancer vaccination. Apart from chimeric antigen receptor (CAR) T cells, which typically recognize antigens in an MHC-independent manner, immunotherapies that engage cytotoxic CD8^+^ T cells depend on the presentation of antigens via MHC class I on the target cell surface. Downregulation or loss of MHC class I on tumor cells has been recognized as a major mechanism of immune evasion (*1–4*). One cancer that has been reported to express low levels of MHC class I is medulloblastoma (*5–7*), the most common malignant brain tumor in children.

Medulloblastoma arises in the cerebellum of infants, children and young adults and has a 5-year survival rate of 73% (*8*). Harsh chemo- and radiotherapy leave survivors with severe long-term side effects, underscoring the need for less toxic and more effective therapies. While immune checkpoint inhibitors have shown promising outcomes in various cancer types, their efficacy in treating medulloblastoma has been limited (*9*). One possible reason for this is the lack of MHC Class I expression on tumor cells, which makes them poor targets for cytotoxic T cells.

Among the molecular subtypes of medulloblastoma, Group 3 tumors, often associated with MYC overexpression or amplification, tend to have the poorest outcomes. Previously, we developed a genetically engineered mouse model that resembles human Group 3 medulloblastoma (*10*). Here, we use this model to study the immune landscape of medulloblastoma and to identify novel approaches to stimulate anti-tumor immune responses. Similar to many human medulloblastoma tumors, our mouse model expresses little MHC class I on the cell surface. Nonetheless, depletion of CD8^+^ T cells accelerates tumor growth, indicating that these cells play an important role in tumor control. We show that T cell-derived interferon gamma (IFNγ) drives MHC class I expression on tumor cells, facilitating T cell killing. To amplify this endogenous anti-tumor response, we combined intratumoral delivery of IFNγ via convection-enhanced delivery (CED) with immune checkpoint blockade. This strategy potentiated T cell–mediated killing and significantly prolonged survival in immunocompetent tumor-bearing mice. These findings highlight the capacity of CD8^+^ T cells to recognize and eliminate medulloblastoma cells when antigen presentation is restored and establish IFNγ as a critical modulator of tumor immunogenicity and a potential adjuvant to T cell–based immunotherapy in medulloblastoma.

## RESULTS

### Medulloblastoma tumors are infiltrated by immune cells

To better understand anti-tumor immune responses in Group 3 medulloblastoma, we analyzed the immune microenvironment in a genetically engineered mouse model of the disease. This model (MP) is based on the orthotopic transplantation of mouse neural stem cells transformed with Myc and a dominant-negative form of p53 (DNp53) (*10*). Importantly, these tumors can grow in syngeneic mice with an intact immune system, making them a valuable model for studying the immune microenvironment. Flow cytometry demonstrated infiltration of tumor tissue by CD45+ immune cells (Fig. 1A, Fig. S1A-D). Constituting more than 55% of all immune cells, myeloid cells were the most abundant immune cell type in the tumor, followed by T cells with 28.4%, with NK cells and B cells present at lower frequencies (11.5% and 6.8% respectively) (Fig. 1A). Notably, CD8^+^ T cells comprised more than 10% of non-tumor cells, highlighting a substantial T cell presence within medulloblastoma tumors.

**Fig. 1.**
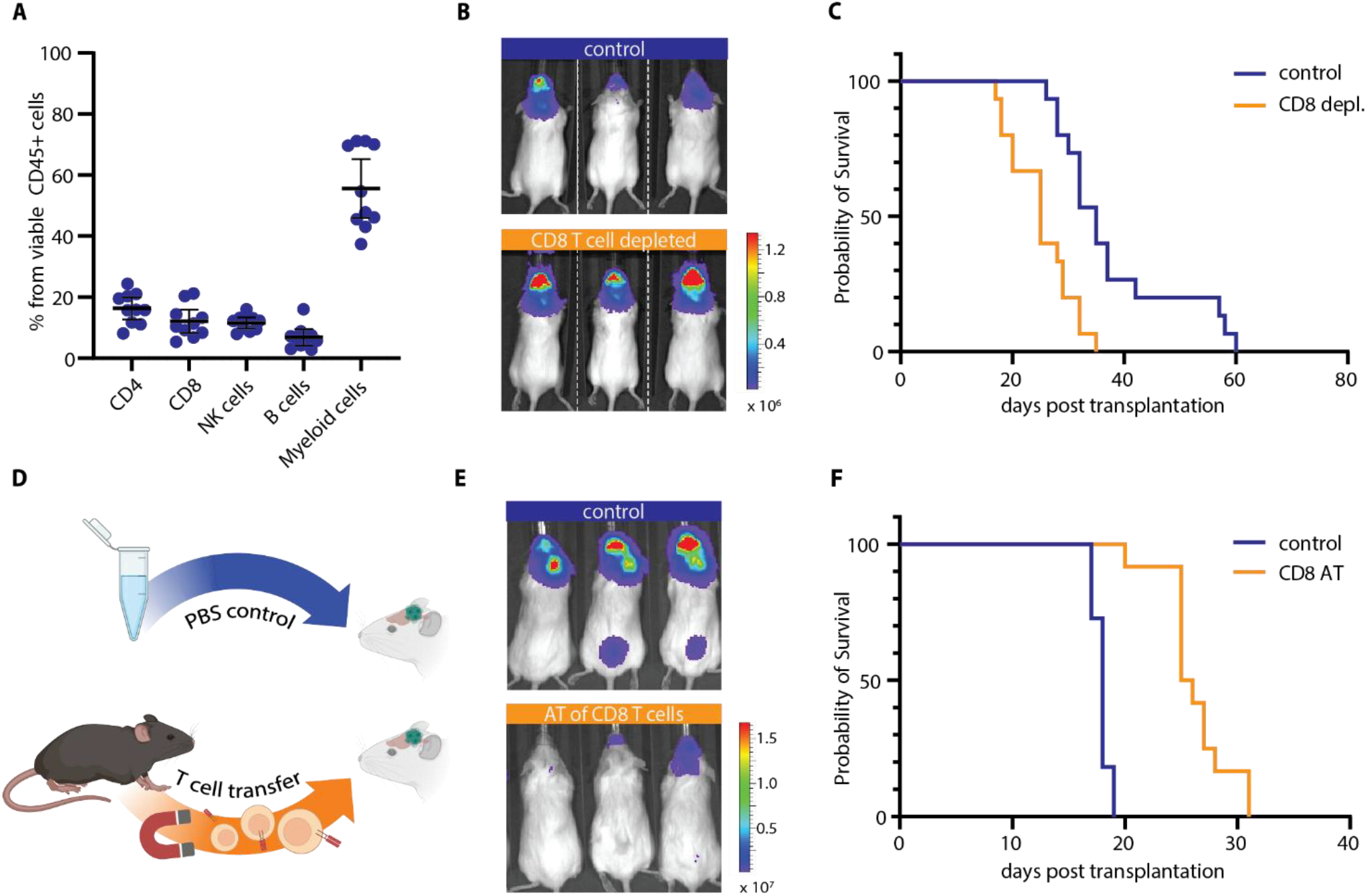
Medulloblastomas are infiltrated and controlled by CD8^+^ T cells. **(A)**Flow cytometry of viable tumor infiltrating immune cells isolated 14d post transplantation, n = 10; mean and 95%CI.**(B)** Representative bioluminescent images of immunocompetent albino C57BL/6 mice injected with mouse medulloblastoma cells. Control animals were treated weekly with an isotype control antibody, CD8^+^ T cell-depleted animals received a CD8-targeting antibody. **(C)** Corresponding survival curve (p = 0.0003; n = 15; log-rank test). **(D)**Schematic representation of adoptive transfer of CD8^+^ T cells isolated from OT-I spleens. **(E)** Representative bioluminescent images of immune-deficient NSG mice injected with OVA positive tumor cells. Experimental animals received adoptive transfer (AT) of 7 – 10 × 10^6 OT-I CD8^+^ T cells (i.v.) 7 – 10 days post tumor transplantation. **(F)**Corresponding survival curve (p = 0.00000094; n(control) = 11; n(CD8 AT) = 12; log-rank test). Bioluminescence scale given in radiance (p/sec/cm^2/sr).

### CD8^+^ T cells control medulloblastoma tumor growth

To evaluate the importance of CD8^+^ T cells in our model, we depleted these cells using anti-CD8 antibodies (Fig. S1E, F) and assessed the effects on tumor growth. Loss of CD8^+^ T cells in otherwise immunocompetent animals resulted in enhanced tumor growth and significantly reduced tumor latency (Fig. 1B, Fig. 1C). Whereas median survival time for control mice was 35 days, CD8^+^ T cell depletion resulted in a median survival of 25 days, suggesting that CD8^+^ T cells play an important role in controlling tumor growth. To assess whether CD8^+^ T cells are sufficient to inhibit tumor growth, we generated tumor cells expressing ovalbumin (OVA) and tested their sensitivity to CD8^+^ T cells from OT-I mice, which express a T cell receptor that recognizes OVA peptide presented by MHC class I (H-2Kb) (*11*). We transferred CD8^+^ T cells isolated from spleens of OT-I mice into the blood of immune-deficient tumor-bearing mice (Fig. 1D). As shown in Figures 1E and 1F, adoptive transfer (AT) of T cells inhibited tumor growth and prolonged survival. These results show that CD8^+^ T cells are functional and capable of controlling tumor growth.

Having observed that CD8^+^ T cells are capable of attacking medulloblastoma tumors, we investigated why they are unable to fully eliminate them. Flow cytometric analysis of tumor cells freshly isolated from mice revealed that the majority of tumor cells lack MHC class I on their surface (Fig. 2A). Consistent with this, flow cytometric analysis of human pediatric brain tumors obtained from surgical resection demonstrated that human medulloblastoma tumors harbored fewer HLA class I-positive tumor cells compared to other pediatric brain tumors, such as high-grade glioma. Medulloblastoma samples also showed lower fluorescence intensity indicative of lower HLA class I expression levels (Fig. 2B, Fig. S2A). Together, these results suggest that medulloblastoma cells may evade CD8^+^ T cell-mediated attack through downregulation of MHC class I.

**Fig. 2.**
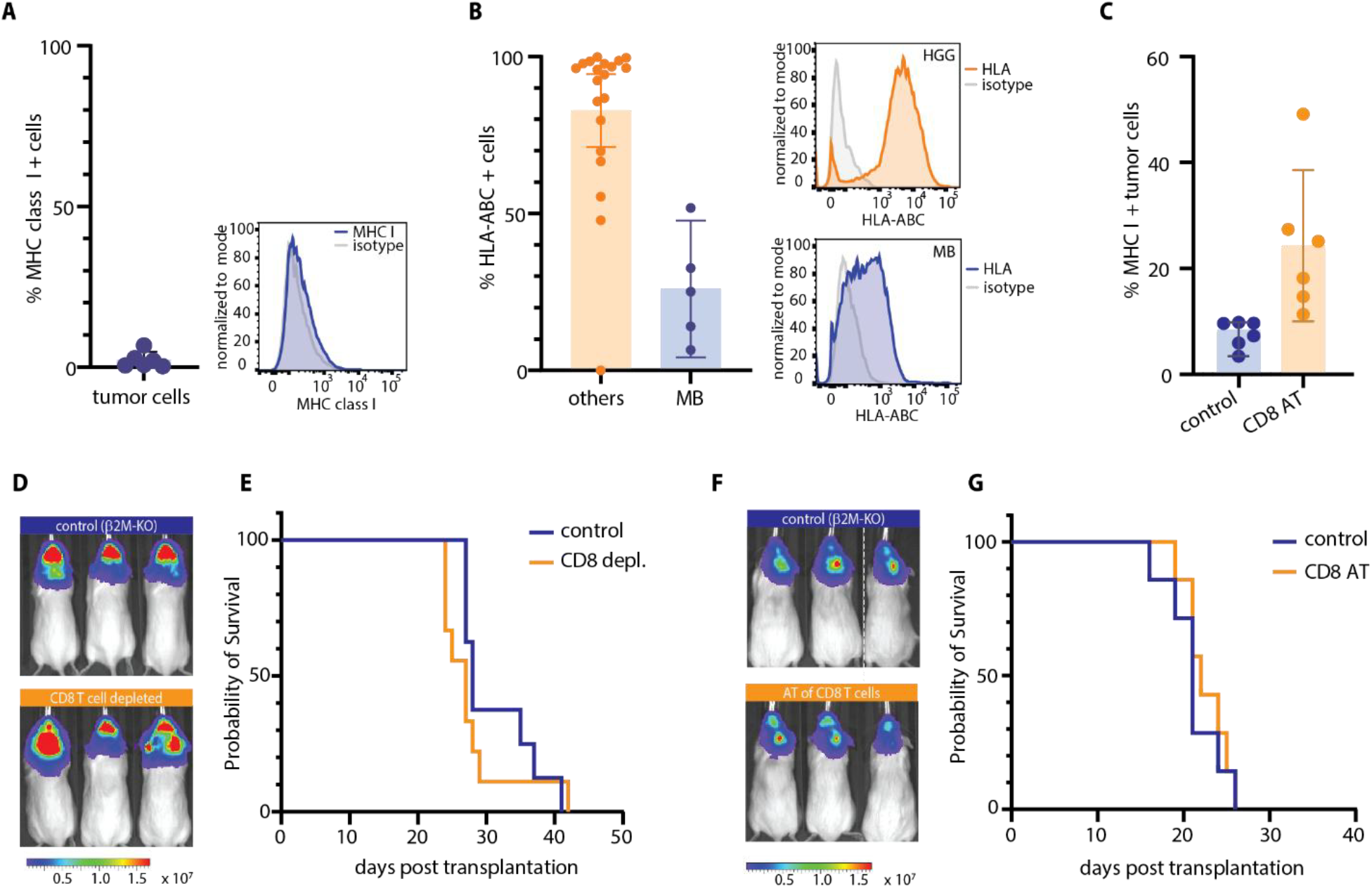
Medulloblastomas evade CD8^+^ T cells by presenting little MHC class I on their surface. **(A)**Flow cytometry of mouse medulloblastoma cells shows a minority of MHC class I positive cells, n = 6; mean and95% CI; and representative histogram of MHC class I staining. **(B)** Flow cytometry of pediatric brain tumors.Medulloblastoma (MB) tumors have fewer cells expressing MHC class I (HLA-ABC) compared to other pediatricbrain cancers, mean and 95% CI, p = 0.0011; Mann Whitney test, n (MB) = 5; n (others) = 20. Representativehistograms of HLA-ABC staining of a high-grade glioma (HGG) and medulloblastoma (MB). **(C)** MHC class Iassessment by flow cytometry of OVA positive tumor cells isolated from NSG mice 3 days after adoptive transfer ofOT-I CD8^+^ T cells, n = 6; p = 0.029, Welch’s t-test, mean and 95% CI. **(D)** Bioluminescent images ofimmunocompetent albino C57BL/6 mice injected with ?2M-KO tumor cells. Control animals were treated weeklywith an isotype control antibody, CD8^+^ T cell-depleted animals received a CD8-targeting antibody. **(E)** Survival curve(p = 0.4088; n = 9, log-rank test). **(F)** bioluminescent images of immune-deficient NSG mice injected with OVA7 positive tumor cells. Adoptive transfer (AT) of OT-I CD8^+^ T cells (i.v.) and **(G)** corresponding survival curve (p = 0.5769; n = 7; log-rank test). Bioluminescence scale given in radiance (p/sec/cm^2/sr).

Although our medulloblastoma cells are largely MHC class I negative ex vivo, which would predict resistance to CD8^+^ T cell–mediated killing, depletion of CD8^+^ T cells accelerates tumor growth. This prompted us to investigate how CD8^+^ T cells can nonetheless restrain tumor growth in vivo. We hypothesized that surface MHC class I expression might be restored at some point in vivo. To test this, we repeated our adoptive transfers of OT-I CD8^+^ T cells into NSG mice bearing OVA positive tumors, but examined MHC expression at earlier time points. Three days after T cell infusion, we found significantly increased numbers of MHC class I positive tumor cells following adoptive transfer of T cells (Fig. 2C, Fig. S2B), suggesting that CD8^+^ T cells may have induced MHC class I expression by tumor cells. To understand whether expression of MHC class I was required for T cell-mediated killing, we generated *beta 2*-*microglobulin*-knockout (β2M-KO) tumor cells. β2M encodes the light chain that together with the alpha heavy chain constitutes the MHC class I complex, and in the absence of β2M, MHC class I cannot be expressed on the surface (*12*). Repeating the CD8^+^ T cell depletion and adoptive transfer experiments with β2M-KO tumors, we did not observe significant differences in tumor burden or survival when depleting or adoptively transferring T cells in these mice (Fig. 2D-G). Taken together, these results demonstrate that CD8^+^ T cells can induce MHC class I expression on tumor cells, and require MHC class I expression to control tumor growth.

### Interferon gamma restores MHC class I expression on medulloblastoma cells

Having demonstrated that T cells can induce MHC class I on tumor cells, we sought to understand the mechanism of this induction. T cells can secrete IFNγ, a potent stimulus for MHC class I expression and antigen presentation (*1, 13, 14*). We thus hypothesized that adoptively transferred OT-I T cells might induce MHC class I expression by secreting IFNγ in the vicinity of tumor cells. Indeed, using an IFNγ reporter mouse strain in which YFP is expressed under the control of the IFNγ promoter, a large proportion of tumor-infiltrating CD8^+^ T cells showed YFP expression by flow cytometry, indicating active IFNγ production (Fig. S3A).

We next analyzed whether immune cell-derived IFNγ induces a response in medulloblastoma cells in vivo. Using cyclic immunofluorescence (cycIF), a high-dimensional imaging technique that enables repeated staining and imaging of the same tissue section, we analyzed tumor sections from control NSG mice and those receiving OT-I CD8^+^ T cells. Mice receiving CD8^+^ T cells showed T cell infiltration into the tumor tissue, and these cells showed cytotoxic activity based on Granzyme B expression (Fig. 3A, 3B). Following adoptive T cell transfer, tumor cells exhibited positive staining for interferon regulatory factor 1 (IRF1) (Fig. 3A, 3D), a well-established marker of IFNγ signaling (*15*), suggesting that CD8^+^ T cells induce interferon signaling in tumor tissue. Our findings suggest that infiltrating CD8^+^ T cells secrete IFNγ, triggering a detectable response in medulloblastoma cells.

**Fig. 3.**
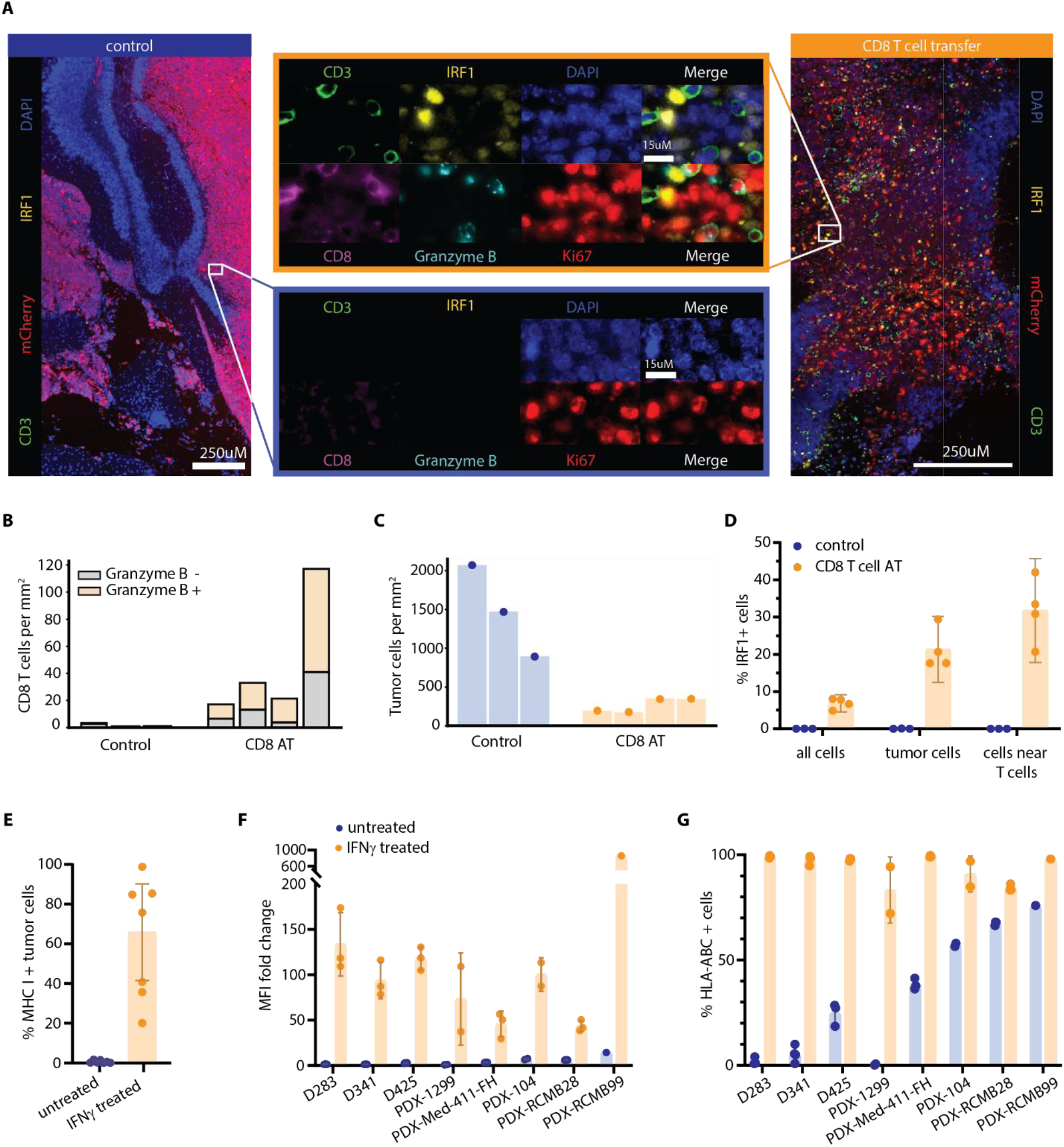
Tumor infiltrating CD8^+^ T cells induce interferon signaling in tumor cells. **(A)**Cyclic immunofluorescent (cycIF) images from immune deficient control mice (n = 3) and mice which have received CD8^+^ T cells (n = 4; AT), and corresponding quantifications of **(B)** infiltrated T cells, **(C)** tumor cells and **(D)**IRF1 positive cells, mean and 95% CI, p = 0.0094; multiple two-sample t-tests, p-values were adjusted for multipletesting using false discovery rate (FDR) correction according to Benjamini and Yekutieli. Tumor cells in AT mice were more than 890 times more likely to express IRF1 than tumor cells in immune deficient mice, Fisher’s exact test, p <0.01. Images were manually gated with the same cutoffs applied across all conditions. Quantification of flowcytometric analysis of in vitro treated **(E)** mouse tumor cells, 10ng/ml mouse IFNγ for 72h, n = 7; p = 0.0156; Wilcoxonsinged-rank test and **(F, G)** human medulloblastoma cells, 100ng/ml human IFNγ 72h, mean and 95% CI. **(F)** Increase in surface HLA-ABC upon IFNγ treatment indicated by mean fluorescence intensity (MFI) fold change compared to isotype control staining and **(G)** corresponding quantification of HLA-ABC positive cells in percentage, n = 3; n(PDX-9 1299 and PDX-104) = 2 and n(RCMB99) = 1; p (D283) = 0.00039; p (D341) = 0.0013; p (D425) = 0.00372; p (Med-411-FH) = 0.0014; p (RCMB28) = 0.0013, PDX-1299; −104 and −RCMB99 n/a. multiple two-sample t-tests, p-values were adjusted for multiple testing using FDR correction according to Benjamini and Yekutieli.

To analyze whether IFNγ can restore MHC class I expression in medulloblastoma cells, we treated tumor cells with IFNγ in vitro and performed flow cytometry. We found a notable upregulation of MHC class I in tumor cells, although the percentage of responding cells varied between tumors (Fig. 3E). We also studied the effect of IFNγ on human medulloblastoma cells. To this end, we treated three cell lines and five patient-derived xenografts (PDXs) in vitro with human IFNγ. We found that the cytokine induced a strong upregulation of MHC class I on the surface of nearly all cells, irrespective of the initial MHC class I levels (Fig. 3F, 3G).

### IFNγ response by tumor cells is essential for CD8^+^ T cell attack

To analyze whether IFNγ treatment sensitizes mouse medulloblastoma cells to CD8^+^ T cell killing, we performed ex vivo co-cultures and real time imaging of cancer cell survival. When OVA-positive tumor cells were co-cultured with OT-I CD8^+^ T cells, continuous cell growth was observed. In contrast, when tumor cells were pre-treated with IFNγ before adding OT-I CD8^+^ T cells, a significantly reduced number of tumor cells was recorded over time compared to tumor cells cultured alone (Fig. 4A). These results demonstrate that IFNγ treatment makes MHC class I negative medulloblastoma cells susceptible to CD8^+^ T cell killing.

**Fig. 4.**
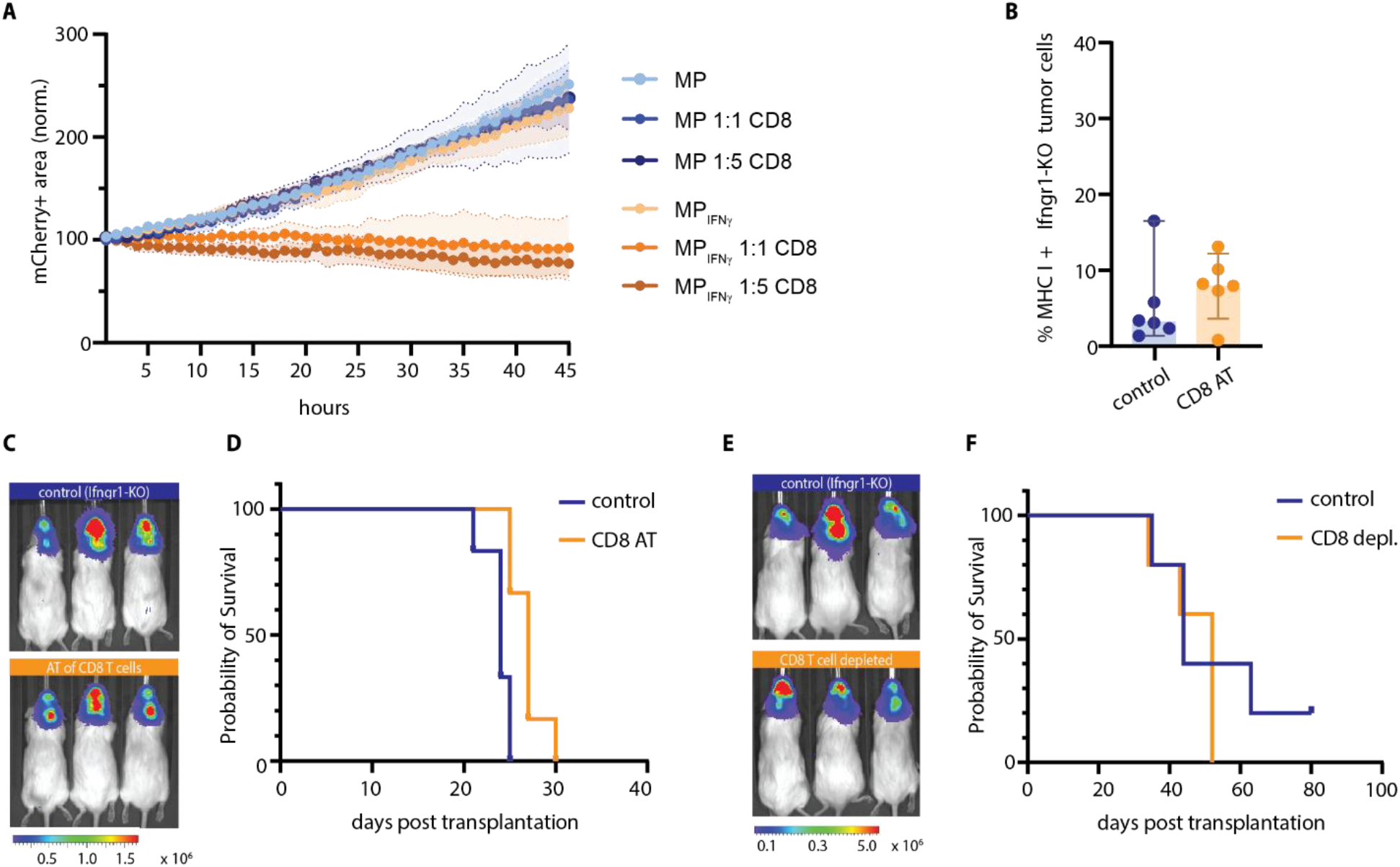
Interferon gamma signaling in tumor cells is required for CD8^+^ T cell attack. **(A)**Live cell imaging of OVA and mCherry positive tumor cells in single or co-culture with OT-I CD8^+^ T cells indifferent effector to target ratios. Tumor cells were either untreated (MP) or pre-treated with 10ng/ml IFNγ for 48h(MPIFNγ). mCherry positive area was quantified and normalized to timepoint 0; mean (thick symbols) and standarderror of the mean (shaded error band) are shown, n = 3. Growth curves were analyzed by repeated-measures two-wayANOVA, with Benjamini–Hochberg correction applied to pairwise group comparisons at each timepoint. Compared to single tumor cell cultures, MPIFNγ tumor cells co-cultured with T cells showed significantly reduced growth from14 h onward at a 1:1 ratio (p < 0.037) and from 11 h onward at a 1:5 ratio (p < 0.035). **(B)** Quantification of MHC class I positive Ifngr1-KO tumor cells by flow cytometry. OVA expressing tumor cells were isolated from NSG mice3 days after adoptive transfer of OT-I CD8^+^ T cells, n = 6; p = 0.31; Mann-Whitney test, mean and 95% CI. **(C)**Representative bioluminescent images of NSG mice injected with OVA positive Ifngr1-KO tumor cells, infused with OT-I CD8^+^ T cells (AT). **(D)** Corresponding survival curve (p = 0.0055; n = 6, log-rank test). **(E)** Representative bioluminescent images of albino C57BL/6 mice injected with Ifngr1-KO tumor cells and either treated with isotypecontrol or anti-CD8 antibody and **(F)** survival curve (p = 0.4655; n = 5, log-rank test). Bioluminescence scale given in radiance (p/sec/cm^2/sr).

Building on our findings that T cells can kill tumor cells exposed to IFNγ ex vivo, we investigated whether the cytokine enhances antigen presentation and T cell-mediated tumor cell killing in vivo. To this end, we treated tumor-bearing NSG mice with an IFNγ sequestering antibody and infused OT-I CD8^+^ T cells. The previously observed upregulation of MHC class I levels following T cell transfer (Fig. 2C) was abrogated by interferon blockade, and no effect of T cells on tumor growth or survival was detected (Fig. S4A, B).

To test whether the lack of interferon was affecting tumor cell susceptibility to T cell killing as opposed to impeding T cell activity, we performed the adoptive transfer experiment with OVA expressing tumor cells that lacked IFNγ receptors (Ifngr1-KO tumor cells). When we isolated tumor cells three days after adoptive transfer of OT-I CD8^+^ T cells, flow cytometry did not show an increase in MHC class I positive tumor cells (Fig. 4B). In line with this result, we only observed a minor effect of CD8^+^ T cell transfer on tumor burden and survival (Fig. 4C, 4D). Strikingly, CD8^+^ T cell depletion in immunocompetent mice that carry Ifngr1-KO tumors had no effect on survival and tumor growth (Fig. 4E, F), indicating that tumor cells that are not responsive to IFNγ remain resistant to CD8^+^ T cell attack. Together these results suggest that T cells secrete IFNγ which induces MHC class I expression in tumor cells, and in turn renders these tumor cells susceptible to CD8^+^ T cell killing.

### IFNγ and anti-PD1 treatment enhance CD8^+^ T cell killing and prolong survival

As our results indicated that IFNγ enables tumor cell killing by cytotoxic CD8^+^ T cells, we hypothesized that increasing the concentration of IFNγ in the tumor might enhance tumor control by CD8^+^ T cells. However, brain tumors are notoriously difficult to treat as many compounds including cytokines do not efficiently penetrate the blood-brain barrier (BBB). Pilot experiments with systemic high-dose IFNγ administration demonstrated limited efficacy in upregulating MHC class I expression on orthotopically transplanted tumor cells (Fig. S5A). Hence, we decided to perform a proof-of-principle experiment by engineering tumor cells to express IFNγ following exposure to doxycycline (which is brain-penetrant). Tumor cells transduced with a construct encoding doxycycline-inducible murine IFNγ (MP-TetOne) responded to doxycycline by increasing expression of MHC class I in vitro (Fig. S5B). OVA-positive MP-OVA-TetOne cells were transplanted into Rag2KO mice for adoptive transfer experiments. Rag2KO animals lack T and B cells but have functional myeloid cells that share the same genetic background (C57BL/6) as transplanted tumor cells and transferred OT-I CD8^+^ T cells. Once tumor growth was verified by bioluminescence imaging, the mice were divided into four groups. Two control groups remained on normal chow, and two experimental groups received doxycycline-containing chow. Two days later, one control group and one doxycycline group received intravenous infusions of OT-I CD8^+^ T cells. As IFNγ is known to induce the expression of the checkpoint protein PD-L1 in tumor cells and innate immune cells (*16–18*), we treated all CD8^+^ T cell infused mice with a weekly i.p. dose of anti-PD1 antibody. The remaining groups were treated with an isotype control antibody. Tumor growth was monitored by bioluminescence imaging. Similar to our previous AT experiments in NSG mice, RagKO mice infused with OT-I CD8^+^ T cells and anti-PD1 antibody showed a delay in tumor growth that resulted in significantly prolonged survival (Fig. 5A, 5B). Tumor growth was further delayed and survival was significantly extended in animals that in addition to CD8^+^ T cells and anti-PD1 received doxycycline chow, mimicking IFNγ treatment (Fig. 5A, 5B).

**Fig. 5.**
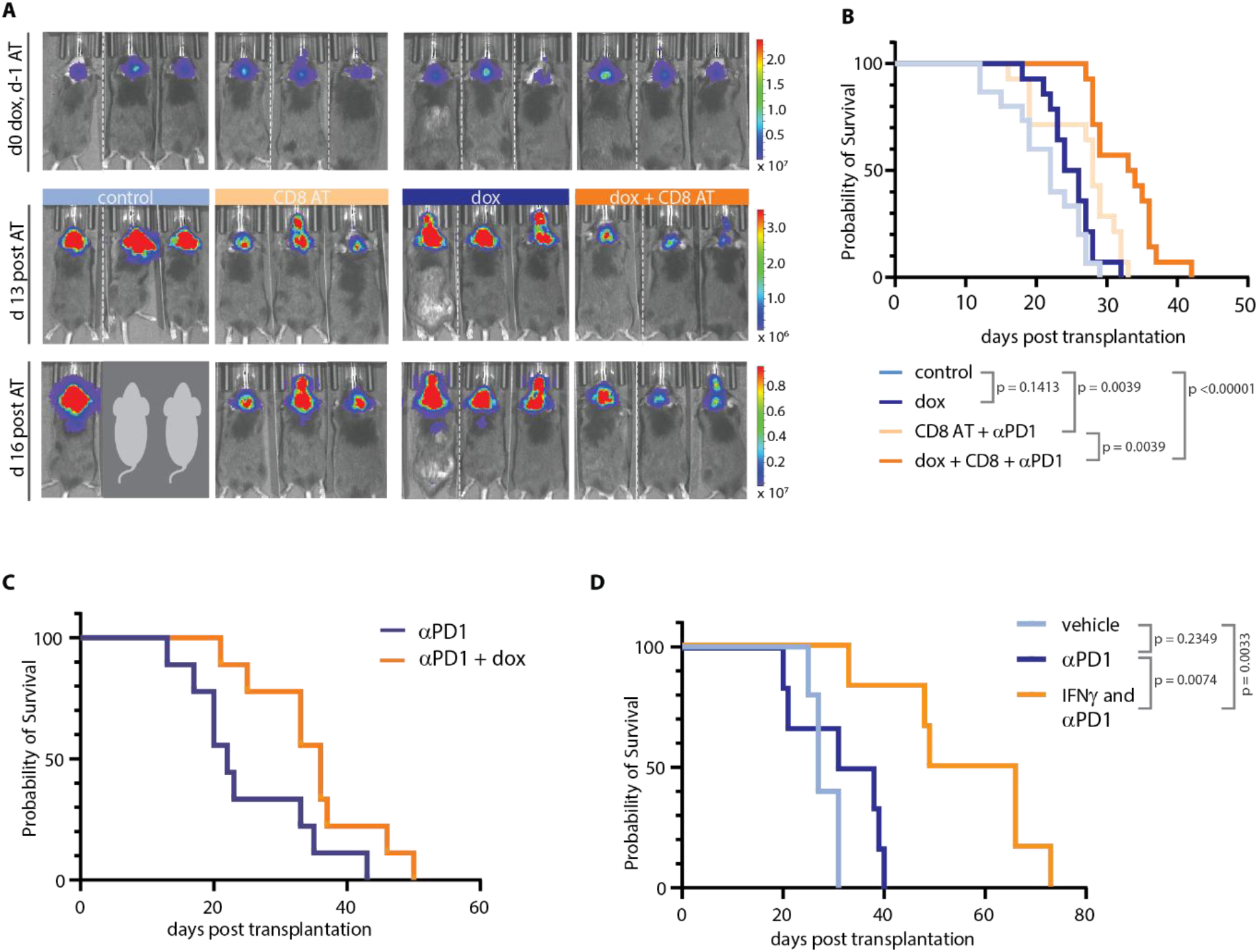
Interferon gamma and anti-PD1 treatment enhance CD8^+^ T cell killing and prolong survival. **(A)**Bioluminescent images of Rag2KO mice injected with MP-OVA-TetOne tumor cells prior to treatment, 13 and16 days post adoptive T cell transfer (AT). Light blue group received control chow and isotype control antibody, darkblue group was fed with doxycycline-containing chow (dox, starting one day before AT) and injected with isotypecontrol antibody, light orange group was subjected to AT of OT-I CD8^+^ T cells and anti-PD-1 treatment; and darkorange group received AT, anti-PD-1 and dox. **(B)** Corresponding survival curves, n = 15; log rank test, multiplecomparisons were corrected using the Benjamini–Hochberg method with a FDR of 5%. **(C)** Kaplan–Meier survival curve of albino C57BL/6 mice injected with MP-TetOne tumor cells, n = 9; p = 0.037; log rank test. Mice were injected weekly with anti-PD1 antibody and received control (blue) or doxycycline (dox) chow (orange). **(D)** Kaplan–Meier survival curves of albino C57BL/6 mice bearing MP tumors and treated via convection-enhanced delivery (CED) with vehicle (n = 5), or IFNγ combined with intraperitoneal (i.p.) anti-PD1 (n = 6), or with anti-PD1 alone (n = 6), log-rank test, multiple comparisons were corrected using the Benjamini–Hochberg method with a FDR of 5%

We next transplanted MP-TetOne tumor cells into immunocompetent albino C57BL/6 mice. After bioluminescence imaging confirmed engraftment of tumors, we divided the mice into a control group and an experimental group, the latter receiving doxycycline food. Both groups received anti-PD1 antibody. Mice fed doxycycline exhibited a median survival time of 36 days compared to 22-day median survival for mice on control chow, suggesting that ectopic IFNγ significantly prolonged survival (Fig. 5C). The results above show that ectopic IFNγ expression by tumor cells enhances CD8^+^ T cell mediated killing and supports their anti-tumor activity. This further suggests a potential clinical value in combining IFNγ with immunotherapy.

To bypass the BBB and implement a clinically relevant therapeutic strategy, we next administered IFNγ to medulloblastoma-bearing mice via convection-enhanced delivery (CED). CED enables direct intratumoral infusion of therapeutics through surgically implanted catheters using a continuous, osmotic pump-driven flow (*19, 20*).

We first tested this approach in immunodeficient NSG mice to determine whether direct intratumoral delivery of IFNγ induces MHC class I upregulation in vivo. Medulloblastoma cells were orthotopically transplanted and tumor growth was confirmed by bioluminescence imaging one week later. Mice were then randomized into vehicle (PBS) and IFNγ treatment groups. A second surgery was performed to position a cannula using the same burr hole created during tumor implantation. Micro-osmotic pump reservoirs containing either IFNγ or vehicle were placed subcutaneously. After three days of treatment, tumors were harvested and dissociated for flow cytometric analysis of MHC class I expression. Results confirmed upregulation of MHC class I in tumor cells (Fig. S5C, D), comparable to levels observed following in vitro treatment with IFNγ.

Having validated that CED of IFNγ effectively enhances MHC class I expression in vivo without inducing noticeable toxicity, we next evaluated its impact on disease progression and overall survival in immunocompetent mice. Medulloblastoma cells were transplanted into the cerebellum, and one week later, mice were stratified into three groups based on equivalent tumor sizes: (i) vehicle (CED) and isotype control antibody (i.p.), (ii) anti-PD1 antibody (i.p.), and (3) combined IFNγ (CED) and anti-PD1 (i.p.). Vehicle and IFNγ were administered for two weeks via ALZET micro-osmotic pumps, while anti-PD1 antibodies were injected weekly throughout the experiment. Survival analysis revealed a median survival of 27 days in the vehicle control group, a non-significant increase to 34.5 days in the anti-PD1 group, and a significantly extended median survival of 57.5 days in mice receiving the combination of IFNγ and anti-PD1 (Fig. 5D).

Together, these results demonstrate that locoregional delivery of IFNγ, in combination with immune checkpoint blockade, significantly improves survival in medulloblastoma-bearing mice.

## DISCUSSION

In this study, we demonstrate the importance of CD8^+^ T cells in controlling medulloblastoma growth and identify IFNγ as an adjuvant for CD8^+^ T cell-centered immunotherapy, which has previously shown limited success as a monotherapy for medulloblastoma.

Medulloblastoma is considered an immunologically cold tumor due to limited immune infiltration and antigen presentation (*21*). Yet, CD8^+^ T cell depletion in our Group 3 medulloblastoma mouse model led to significantly increased tumor progression, and complementary adoptive transfer experiments of CD8^+^ T cells in immune-deficient tumor bearing mice resulted in significantly prolonged survival. These results demonstrate that even a small number of CD8^+^ T cells can counteract medulloblastoma growth despite minimal MHC class I expression on tumor cells. Supporting this, emerging evidence suggests that medulloblastoma elicits T cell activation in patients. Previous studies identified neoantigens capable of triggering T cell activation (*22*), and single-cell RNA sequencing combined with T cell receptor profiling revealed clonal expansion of cytotoxic CD8^+^ T cells within the medulloblastoma microenvironment (*23*). Together, these findings underscore the presence of tumor-reactive CD8^+^ T cells in medulloblastoma and highlight the need for strategies to amplify these responses to achieve durable tumor rejection.

Our results suggest that low MHC class I expression represents a key barrier to effective CD8^+^ T cell-responses in medulloblastoma. Notably, we demonstrate that exposure to IFNγ induces expression of surface MHC class I on medulloblastoma cells, rendering them more susceptible to cytotoxic CD8^+^ T cell attack. This is in line with a recent report showing that clinical response to immunotherapy in melanoma corresponded most highly with an IFNγ signature in tumor cells, suggesting that interferon signaling may serve to jump-start anti-tumor immune responses (*24*). Of note, IFNγ may further exert its therapeutic benefit by reprogramming myeloid cells toward a pro-inflammatory phenotype (*25, 26*). The cytokine has also been reported to enhance cytotoxicity of T cells and natural killer cells (*26, 27*). Current research is leveraging these immune modulatory and pro-inflammatory properties to enhance anti-tumor immunity (*28, 29*), as the direct cytotoxic effects of IFNγ on tumor cells require high, toxic doses with potentially limited therapeutic benefit. Despite its potential, IFNγ is not currently approved for cancer immunotherapy, and its systemic delivery is associated with challenges including a short half-life, rapid absorption by tissues, and a dependency on precise dosing schedules. IFNγ can also induce counter-regulatory pathways such as PD-L1 upregulation (*28–30*), which we found particularly relevant in medulloblastoma where infiltrating CD8^+^ T cells display substantial PD-1 expression (*31*). This provides strong rationale for combining IFNγ–mediated sensitization with PD-1 checkpoint blockade. Moreover, the BBB significantly limits the passive diffusion of large molecules like cytokines. With studies suggesting that only about 0.1% of plasma interferon alpha reaches the brain following systemic delivery (*32*), IFNγ is likely to exhibit similarly poor permeability.

To circumvent these limitations, we initially employed an inducible vector for ectopic IFNγ expression within tumor cells. Upon induction, this strategy resulted in enhanced antigen presentation and a significant survival benefit, supporting the therapeutic potential of IFNγ in this context. Building on this insight, we pursued a more translatable treatment approach, CED, to administer IFNγ directly into brain tumors via surgically implanted osmotic pumps. CED is still an experimental technique, but recent clinical trials have demonstrated its promise for delivery of therapeutic agents to brain cancer patients (*33*). Here we show that CED of IFNγ not only induced MHC class I expression in vivo but in combination with anti-PD1 checkpoint blockade also markedly prolonged survival in medulloblastoma-bearing mice. These findings validate CED as a clinically relevant delivery method for IFNγ that can effectively sensitize immunologically cold tumors to CD8^+^ T cell–mediated killing. Future studies will need to address potential challenges such as resistance to IFNγ signaling (*34*), alternative immune evasion mechanisms (*35*), and dose-related toxicities, which will be critical to fully harness its therapeutic potential.

In summary, we present novel findings demonstrating that medulloblastoma responds to IFNγ treatment with augmented antigen presentation via MHC class I and increased susceptibility to CD8^+^ T cell-mediated killing. Given the low immunogenicity of medulloblastoma, our finding that IFNγ treatment can increase tumor visibility to the immune system is particularly significant. By integrating mechanistic insights with translatable delivery strategies, our work highlights IFNγ as a promising agent to sensitize immunologically cold brain tumors to T cell therapies.

## MATERIALS AND METHODS

### Study design

This study aimed to investigate the role of CD8^+^ T cells in the anti-medulloblastoma immune response using a combination of in vitro and in vivo approaches. We employed a mouse model of medulloblastoma, using syngeneic immunocompetent albino C57BL/6J mice and immune-deficient NSG or RagKO mice, in conjunction with ovalbumin (OVA)-expressing tumor cells and CD8^+^ T cells isolated from OT-I transgenic mice. Experimental approaches included CD8^+^ T cell depletion, adoptive cell transfer, and pharmacological interventions using CED to increase intratumoral IFNγ levels and thus sensitize tumor cells to T cell killing.

Animals were stratified into experimental groups based on equivalent tumor burden as measured by bioluminescence imaging. For survival analyses, mice were euthanized upon reaching predefined humane endpoints (including neurological symptoms, signs of distress, or >20% body weight loss).

To validate and extend findings to human disease, we performed flow cytometry on freshly isolated patient material as well as in vitro assays with established human medulloblastoma cell lines and patient-derived xenografts (PDXs), assessing MHC class I upregulation in response to IFNγ treatment.

### Mouse strains

Tumor cells were injected into immunocompetent B6(Cg)-Tyr^c-2J/^J (albino B6; JAX strain #00058); these are C57BL/6 mice that carry a mutation in the tyrosinase gene resulting in complete absence of pigment from skin, hair and eyes, which allows for more accurate quantification of bioluminescence imaging.

Other mouse strains that served as hosts for tumor cell injections are NOD.Cg-Prkdc^scid^ Il2^rgtm1Wjl^/SzJ (NSG, JAX strain #005557), B6.Cg-Rag2^tm1.1Cgn/^J (Rag2KO, JAX strain # 008449) and B6.129S4-Ifng^tm3.1Lky^/J (GREAT, JAX strain # 017581) mice.

Mice used for tumor cell generation include B6(Cg)-Tyr^c-2J/^J; B6.129S7-Ifngr1^tm1Agt^/J (IfngrKO, Jax strain # 003288), and B6.129P2-B2m^tm1Unc^/DcrJ (b2MKO, JAX strain # 002087).

C57BL/6-Tg(TcraTcrb)1100Mjb/J (OT-I, JAX strain # 003831) mice were used to isolate CD8^+^ T cells.

This study was approved by the animal care and use committees at Sanford Burnham Prebys, the University of California San Diego (UCSD) and Columbia University. All experiments were performed in accordance with national guidelines and regulations for the care and use of laboratory animals.

### Mouse tumor generation

CD133+ neural stem cells from cerebella of newborn mice were isolated and infected with Myc^T58A^-IRES-luciferase and DNp53-IRES-mCherry retroviruses, followed by stereotactic injection into the cerebellum of adult mice (*10*). Developed tumors were harvested and re-transplanted for experiments.

To generate OVA-expressing tumors, tumor cells were transduced with lentivirus encoding chicken ovalbumin. For this, Addgene plasmid #113030 was modified to encode ovalbumin-P2A-GFP. To create tumors with doxycycline-inducible overexpression of murine IFNγ, tumor cells were transduced with retrovirus encoding *Ifng* cloned into pRetroX-TetOne-Puro vector by Takara/Clontech.

### Intracranial injections

Anesthetized mice were positioned in a stereotaxic frame, and a midline incision was made to expose the skull. Tumor cells (2 × 10^4^ mouse or 2.5 × 10^5^ human PDX cells in 3 µL of sterile PBS) were aspirated into a Hamilton syringe fitted with a 27½-gauge needle. The needle was carefully advanced to a depth of 1.5 mm - 2 mm, approximately 1 mm lateral to the midline and 1mm posterior to the lambda suture.

For tumor dissociation, tissue was incubated in 10 U/ml papain (Worthington Biochemical) and DNase solution followed by gentle mechanical dissociation, unless cells were subjected to flow cytometry. For these assays, tumors were dissociated using 175μg/ml Liberase (Sigma-Aldrich) and DNase in PBS.

### Patient material

Human tumors were obtained from Rady Children’s Hospital after surgical tumor dissection. Samples included 5 medulloblastomas (MB), 10 high-grade gliomas (HGG), 2 diffuse midline gliomas (DMG), 4 ependymomas (EP), 1 pleomorphic xanthoastrocytoma (PXA), 2 atypical teratoid rhabdoid tumors (ATRT) and 1 central nervous system high-grade neuroepithelial tumor with BCL6 co-repressor (BCOR) internal tandem duplication. Tissues were dissociated using the above-mentioned Liberase/DNase-based protocol.

The studies involving human samples were approved by the institutional review boards at Sanford Burnham Prebys and UCSD and conducted in accordance with the Declaration of Helsinki. All participants provided written informed consent.

### Bioluminescence imaging

Tumor progression was monitored using bioluminescence imaging in mice engrafted with luciferase-expressing MP cells. Mice were injected i.p. with D-luciferin (PerkinElmer; 75 mg/kg) and anesthetized with isoflurane for imaging. Twelve minutes post-luciferin injection, mice were imaged with exposure times depending on signal intensity. The cranial region was defined as the region of interest (ROI), and total photons were quantified using Living Image Software (PerkinElmer, Waltham, MA, USA). Imaging was performed at regular intervals to assess tumor growth and response to experimental treatments.

### Adoptive transfers

CD8^+^ T cells were isolated from the spleens of OT-I mice (JAX Strain #003831) using the MagniSort™ Mouse CD8^+^ T cell Enrichment Kit (Thermo Fisher Scientific), and cultured in RMPI medium supplemented with 10% FBS, 2mM L-glutamine, 1x non-essential amino acids, 1x sodium pyruvate, 10mM HEPES buffer, P/S, 50μM β-mercaptoethanol, 10ng/ml mouse IL-2 (Biolegend) and 15μl CD3/CD28 activator beads (Thermo Fisher) per 1 × 10^6^ cells per ml.

Cells were cultured for 5-10 days before beads were removed and 7 – 10 × 10^6^ cells were injected in 50μl PBS i.v. (retro-orbitally) into tumor-bearing mice. Control mice received 50μl PBS.

For experiments where anti-PD1 treatment was indicated, 5μg/ml anti-PD1 antibody was added to the T cell suspension before injection into mice. In addition, mice were treated with a weekly i.p. dose of anti-PD1 antibody.

### In vivo treatments

Antibodies used for in vivo experiments were obtained from BioXcell and administered i.p. weekly at a dose of 200μg/mouse. Anti-PD1 antibody dose was lowered to 100μg/mouse after the initial dose. Anti-IFNγ antibody was administered every 3 days.

Systemic IFNγ treatment was performed in NSG mice. 40μg murine IFNγ (Biolegend) was administered i.p. daily over the course of two days.

Doxycycline containing food (1000mg/kg) was purchased from Envigo.

### Convection-enhanced delivery (CED)

One week after intracranial tumor transplantation, CED pumps were implanted in subcutaneous pockets between the shoulder blades of tumor-bearing mice. These micro-osmotic pumps were loaded with 20μg/ml or 80μg/ml to achieve a delivery of 20ng/hour over the course of 3 days (ALZET, #1003D) or 14 days (ALZET, #1002), respectively. A cannula was stereotaxically positioned at the site of prior tumor implantation using a Brain Infusion kit 3 (ALZET, #0008851), adjusted to a depth of 2 mm. The cannula was secured to the skull with Loctite adhesive (ALZET, #0008670) and reinforced with dental cement (Fisher Scientific, #10-000-786). The drug or vehicle (PBS) was delivered at a rate of 1μl/hour for 3 days or 0.25μl/hour for 14 days. At that time, pumps were surgically removed and either tumors dissociated for flow cytometry or mice observed daily for progression of the disease and predefined endpoint criteria. Upon manifestation of neurological or morbidity symptoms, animals were euthanized.

### Cell culture

Mouse tumor cells were cultured as spheroids in Neurobasal-A media (Invitrogen), supplemented with 1X B-27-A (Thermo Fisher Scientific), 1X N-2 (Thermo Fisher Scientific), 2mM L-glutamine, P/S and dissociated with Accutase (Gibco).

Human cell lines (D-283 Med; D-341 Med; D-458; from ATCC) were cultured in DMEM containing 10% FBS, 2mM L-glutamine, P/S. Human PDX were cultured in above described Neurobasal-A medium. Human cells were dissociated with TrypLE (Gibco).

### Live-cell Imaging and Cytotoxicity Assay (Incucyte)

To assess CD8^+^ T cell–mediated cytotoxicity in real time, we used the Incucyte S3 Live-Cell Analysis System (Sartorius). mCherry^+^ medulloblastoma tumor cells were pretreated with 100 ng/mL IFNγ or remained untreated and were seeded the following day into laminin- and poly-L-ornithine–coated 96-well flat-bottom plates at a density of 7.5 × 10^3^ cells per well. After overnight adherence, OT-I CD8^+^ T cells were added at the indicated effector-to-target (E:T) ratios. Co-cultures were maintained in Neurobasal-A medium supplemented with glutamine, B27, N2 (as described above), and 2 ng/mL recombinant murine IL-2 (Biolegend). Plates were imaged every hour for 48 hours, three wells per sample and 2 fields per well. Average mCherry^+^ tumor cell area was quantified based on fluorescence using Incucyte software. mCherry^+^ area over time was normalized to the initial time point (t0). Three independent experiments were performed and cytotoxicity was interpreted as a decrease in normalized mCherry^+^ signal relative to tumor-only control wells.

### Flow cytometry

Human and mouse tumors were dissociated using Liberase solution as described above, stained in 1% FBS/PBS with anti-HLA (Biolegend, clone W6/32), or anti-MHC class I (Invitrogen, clone 34-1-2S), or isotype control (Biolegend, clone MOPC-173).

Cultured cells were dissociated with Accutase and processed as described above for flow cytometry.

Interferon treatments were conducted using 10ng/ml mouse IFNγ purchased from Biolegend, and 100ng/ml human IFNγ purchased from GenScript.

Mouse CD8^+^ T cells were detected using anti-CD8 antibody (Biolegend, clone 53-6.7), anti-CD3ε (Biolegend, clone 17A2), anti-CD45 (Biolegend, clone 30-F11). Other antibodies used include anti-CD11b (Biolegend, clone M1/70), anti-CD4 (Biolegend, clone GK1.5), anti-B220 (Biolegend, clone RA3-6B2) or anti-CD19 (eBioscience, clone ID3), anti-NK1.1 (BD, clone PK136). Zombie Yellow (Biolegend) or DAPI (Sigma-Aldrich) served as viability dyes.

Flow cytometry results were analyzed using FlowJo™ Software, BD Life Sciences.

### Cyclic Immunofluorescence

A set of antibodies was validated on mouse lymphoid and cerebellar tissues: anti-CD3 (Abcam #ab135372; clone SP162), anti-IRF1 (Cell Signaling Technology #14105S; clone D5E4), anti-Ki67 (Cell Signaling Technology #12075; clone D3B5), anti-CD8 (Cell Signaling Technology #98941; clone D4W2Z), anti-GRNZB (Cell Signaling Technology #44153; clone E5V2L), anti-mCherry (Abcam #ab167453; polyclonal). Formalin-fixed, paraffin-embedded brain sections from tumor-bearing mice were deparaffinized, rehydrated, and subjected to antigen retrieval, blocking, and iterative staining with directly conjugated primary antibodies according to Eng et al. (*36, 37*). Regions of interest (ROIs) enriched in tumor cells were selected based on mCherry expression.

Images were registered using the mplexable pipeline (*36*). Nuclear segmentation was performed in CellProfiler (v4.2.4) based on DAPI signal, and marker intensities were quantified in either the nucleus (for IRF1) or a 5 μm expanded perinuclear zone (for surface markers). Thresholding of marker expression was performed using Gaussian Mixture Modeling, followed by manual curation. CD8^+^ T cells were identified by CD45, CD3 and CD8 co-staining. Tumor cells were identified as mCherry^+^CD45^-^ cells.

To assess IRF1 expression relative to T cell localization, all cells within 25 μm of an activated CD8^+^ T cell (CD45^+^CD3^+^CD8^+^GranzymeB^+^) were classified as proximal. Spatial coordinates of cell centroids were derived in CellProfiler and analyzed using Scipy’s KDTree function. A Fisher’s exact test was applied to determine whether tumor cell IRF1 status was independent of treatment condition. Additionally, the odds ratio of IRF1^+^/IRF1^-^ tumor cells was computed between conditions.

### Statistics

Data were plotted and analyzed using GraphPad Prism version 10.0.0 for Windows, GraphPad Software, Boston, Massachusetts USA. Survival was plotted using Kaplan-Meier curves and analyzed with the log-rank test. Survival curve comparisons were adjusted for multiple testing using the Benjamini–Hochberg method, controlling the FDR at 5%. Normality was assessed using the Shapiro–Wilk test. For comparisons between two groups, the following tests were applied:

1. a two-tailed unpaired Student’s t-test for normally distributed, independent samples;
2. multiple paired t-tests for treated vs. untreated samples, with p-values adjusted using the Benjamini–Krieger–Yekutieli two-stage step-up procedure to control the FDR;
3. the Wilcoxon matched-pairs signed-rank test for paired data that did not meet the assumptions of normality;
4. the Mann–Whitney U test for non-normally distributed, unpaired samples.

To analyze longitudinal growth in our live cell imaging experiment, we applied a repeated-measures two-way ANOVA using the *aov_ez* function from the *afex* R package, specifying time as the within-subject factor and group as the between-subject factor. This model tests for overall effects of time, group differences, and potential time × group interactions. To identify at which specific timepoints groups differed, we performed pairwise comparisons between groups at each hour, adjusting p-values for multiple testing with the Benjamini–Hochberg procedure.

## List of Supplementary Materials

Figures S1 – S5 in supplement.

## ACKNOWLEDGEMNTS

We kindly thank the Sanford Burnham Prebys and UCSD Animal Facilities for assistance with colony maintenance and animal care; the Sanford Burnham Prebys Bioinformatics core facility for support with data analysis and statistics; the Sanford Burnham Prebys Flow Cytometry core facilities and the Human Embryonic Stem Cell Core at UCSD for help with FACS sorting. We thank Dr. Anusha Preethi Ganesan for assistance with data analysis. We gratefully acknowledge Rady Children’s Hospital, and particularly Drs. John Crawford, Denise Malicki and Mike Levy for providing patient samples. We thank Bryan Hall for invaluable support with regulatory approvals and logistical coordination. We are grateful to Syber Haverlack for help with CycIF staining, imaging and post image processing. Schematic was created with BioRender.com.

## AUTHOR CONTRIBUTIONS

Conceptualization: RJWR, TE

Methodology: TE, MBM, IY, KC

Investigation: TE, MBM, TT, VP, KRC, IY, KC, AD, LC

Visualization: TE, VP

Funding acquisition: RJWR, EF, JPM, LC, SLP, KC

Project administration: RJWR, TE

Supervision: RJWR

Writing – original draft: RJWR, TE

Writing – review & editing: RJWR, TE

## DATA AND MATERIAL AVAILABILITY

All data are available in the main text or the supplementary materials.

## REFERENCES

1. K. Dhatchinamoorthy, J. D. Colbert, K. L. Rock, Cancer Immune Evasion Through Loss of MHC Class I Antigen Presentation. Front Immunol 12, 636568 (2021).

2. G. Sari, K. L. Rock, Tumor immune evasion through loss of MHC class-I antigen presentation. Curr Opin Immunol 83, 102329 (2023).

3. P. Thor Straten, F. Garrido, Targetless T cells in cancer immunotherapy. J Immunother Cancer 4, 23 (2016).

4. C. Galassi, T. A. Chan, I. Vitale, L. Galluzzi, The hallmarks of cancer immune evasion. Cancer Cell 42, 1825–1863 (2024).

5. D. Haydar, H. Houke, J. Chiang, Z. Yi, Z. Odé, K. Caldwell, X. Zhu, K. S. Mercer, J. L. Stripay, T. I. Shaw, P. Vogel, C. DeRenzo, S. J. Baker, M. F. Roussel, S. Gottschalk, G. Krenciute, Cell-surface antigen profiling of pediatric brain tumors: B7-H3 is consistently expressed and can be targeted via local or systemic CAR T-cell delivery. Neuro Oncol 23, 999–1011 (2021).

6. L. Raffaghello, P. Nozza, F. Morandi, M. Camoriano, X. Wang, M. L. Garrè, A. Cama, G. Basso, S. Ferrone, C. Gambini, V. Pistoia, Expression and functional analysis of human leukocyte antigen class I antigen-processing machinery in medulloblastoma. Cancer Res 67, 5471–5478 (2007).

7. J. F. Vermeulen, W. Van Hecke, E. J. M. Adriaansen, M. K. Jansen, R. G. Bouma, J. Villacorta Hidalgo, P. Fisch, R. Broekhuizen, W. G. M. Spliet, M. Kool, N. Bovenschen, Prognostic relevance of tumor-infiltrating lymphocytes and immune checkpoints in pediatric medulloblastoma. Oncoimmunology 7, e1398877 (2018).

8. Q. T. Ostrom, H. Gittleman, G. Truitt, A. Boscia, C. Kruchko, J. S. Barnholtz-Sloan, CBTRUS Statistical Report: Primary Brain and Other Central Nervous System Tumors Diagnosed in the United States in 2011-2015. Neuro Oncol 20, iv1–iv86 (2018).

9. I. J. Dunkel, F. Doz, N. K. Foreman, D. Hargrave, A. Lassaletta, N. André, J. R. Hansford, T. Hassall, M. Eyrich, S. Gururangan, U. Bartels, A. Gajjar, L. Howell, D. Warad, M. Pacius, R. Tam, Y. Wang, L. Zhu, K. Cohen, Nivolumab with or without ipilimumab in pediatric patients with high-grade CNS malignancies: Safety, efficacy, biomarker, and pharmacokinetics-CheckMate 908. Neuro Oncol 25, 1530–1545 (2023).

10. Y. Pei, C. E. Moore, J. Wang, A. K. Tewari, A. Eroshkin, Y. J. Cho, H. Witt, A. Korshunov, T. A. Read, J. L. Sun, E. M. Schmitt, C. R. Miller, A. F. Buckley, R. E. McLendon, T. F. Westbrook, P. A. Northcott, M. D. Taylor, S. M. Pfister, P. G. Febbo, R. J. Wechsler-Reya, An animal model of MYC-driven medulloblastoma. Cancer Cell 21, 155–167 (2012).

11. K. A. Hogquist, S. C. Jameson, W. R. Heath, J. L. Howard, M. J. Bevan, F. R. Carbone, T cell receptor antagonist peptides induce positive selection. Cell 76, 17–27 (1994).

12. B. H. Koller, P. Marrack, J. W. Kappler, O. Smithies, Normal development of mice deficient in beta 2M, MHC class I proteins, and CD8+ T cells. Science 248, 1227–1230 (1990).

13. V. Shankaran, H. Ikeda, A. T. Bruce, J. M. White, P. E. Swanson, L. J. Old, R. D. Schreiber, IFNgamma and lymphocytes prevent primary tumour development and shape tumour immunogenicity. Nature 410, 1107–1111 (2001).

14. J. Wang, Q. Lu, X. Chen, I. Aifantis, Targeting MHC-I inhibitory pathways for cancer immunotherapy. Trends Immunol 45, 177–187 (2024).

15. A. Garcia-Diaz, D. S. Shin, B. H. Moreno, J. Saco, H. Escuin-Ordinas, G. A. Rodriguez, J. M. Zaretsky, L. Sun, W. Hugo, X. Wang, G. Parisi, C. P. Saus, D. Y. Torrejon, T. G. Graeber, B. Comin-Anduix, S. Hu-Lieskovan, R. Damoiseaux, R. S. Lo, A. Ribas, Interferon Receptor Signaling Pathways Regulating PD-L1 and PD-L2 Expression. Cell Rep 19, 1189–1201 (2017).

16. R. D. Dorand, J. Nthale, J. T. Myers, D. S. Barkauskas, S. Avril, S. M. Chirieleison, T. K. Pareek, D. W. Abbott, D. S. Stearns, J. J. Letterio, A. Y. Huang, A. Petrosiute, Cdk5 disruption attenuates tumor PD-L1 expression and promotes antitumor immunity. Science 353, 399–403 (2016).

17. P. Loke, J. P. Allison, PD-L1 and PD-L2 are differentially regulated by Th1 and Th2 cells. Proc Natl Acad Sci U S A 100, 5336–5341 (2003).

18. M. Zibelman, A. W. t. MacFarlane, K. Costello, T. McGowan, J. O’Neill, R. Kokate, H. Borghaei, C. S. Denlinger, E. Dotan, D. M. Geynisman, A. Jain, L. Martin, E. Obeid, K. Devarajan, K. Ruth, R. K. Alpaugh, E. A. Dulaimi, E. Cukierman, M. Einarson, K. S. Campbell, E. R. Plimack, A phase 1 study of nivolumab in combination with interferon-gamma for patients with advanced solid tumors. Nat Commun 14, 4513 (2023).

19. M. Gallitto, X. Zhang, G. De Los Santos, H. J. Wei, E. C. Fernández, S. Duan, G. Sedor, N. Yoh, D. Kokossis, J. C. Angel, Y. F. Wang, E. White, C. J. Kinslow, X. Berg, L. Tomassoni, F. Zandkarimi, I. I. C. Chio, P. Canoll, J. N. Bruce, N. A. Feldstein, R. D. Gartrell, S. K. Cheng, J. H. Garvin, S. Zacharoulis, R. J. Wechsler-Reya, J. Pavisic, A. Califano, Z. Zhang, C. C. Wu, Targeted delivery of napabucasin with radiotherapy improves outcomes in diffuse midline glioma. Neuro Oncol 27, 795–810 (2025).

20. A. M. Sonabend, A. S. Carminucci, B. Amendolara, M. Bansal, R. Leung, L. Lei, R. Realubit, H. Li, C. Karan, J. Yun, C. Showers, R. Rothcock, J. O A. Califano, P. Canoll, J. N. Bruce, Convection-enhanced delivery of etoposide is effective against murine proneural glioblastoma. Neuro Oncol 16, 1210–1219 (2014).

21. T. Eisemann, R. J. Wechsler-Reya, Coming in from the cold: overcoming the hostile immune microenvironment of medulloblastoma. Genes Dev 36, 514–532 (2022).

22. F. Blaeschke, M. C. Paul, M. U. Schuhmann, A. Rabsteyn, C. Schroeder, N. Casadei, J. Matthes, C. Mohr, R. Lotfi, B. Wagner, T. Kaeuferle, J. Feucht, S. Willier, R. Handgretinger, S. StevanoviC, P. Lang, T. Feuchtinger, Low mutational load in pediatric medulloblastoma still translates into neoantigens as targets for specific T-cell immunotherapy. Cytotherapy 21, 973–986 (2019).

23. A. Upadhye, K. E. Meza Landeros, C. Ramírez-Suástegui, B. J. Schmiedel, E. Woo, S. J. Chee, D. Malicki, N. G. Coufal, D. Gonda, M. L. Levy, J. A. Greenbaum, G. Seumois, J. Crawford, W. D. Roberts, S. P. Schoenberger, H. Cheroutre, C. H. Ottensmeier, P. Vijayanand, A. P. Ganesan, Intra-tumoral T cells in pediatric brain tumors display clonal expansion and effector properties. Nat Cancer 5, 791–807 (2024).

24. C. S. Grasso, J. Tsoi, M. Onyshchenko, G. Abril-Rodriguez, P. Ross-Macdonald, M. Wind-Rotolo, A. Champhekar, E. Medina, D. Y. Torrejon, D. S. Shin, P. Tran, Y. J. Kim, C. Puig-Saus, K. Campbell, A. Vega-Crespo, M. Quist, C. Martignier, J. J. Luke, J. D. Wolchok, D. B. Johnson, B. Chmielowski, F. S. Hodi, S. Bhatia, W. Sharfman, W. J. Urba, C. L. Slingluff, Jr., A. Diab, J. Haanen, S. M. Algarra, D. M. Pardoll, V. Anagnostou, S. L. Topalian, V. E. Velculescu, D. E. Speiser, A. Kalbasi, A. Ribas, Conserved Interferon-γ Signaling Drives Clinical Response to Immune Checkpoint Blockade Therapy in Melanoma. Cancer Cell 38, 500–515.e503 (2020).

25. D. Duluc, M. Corvaisier, S. Blanchard, L. Catala, P. Descamps, E. Gamelin, S. Ponsoda, Y. Delneste, M. Hebbar, P. Jeannin, Interferon-gamma reverses the immunosuppressive and protumoral properties and prevents the generation of human tumor-associated macrophages. Int J Cancer 125, 367–373 (2009).

26. J. Remsik, X. Tong, R. Z. Kunes, M. J. Li, R. Estrera, J. Snyder, C. Thomson, A. M. Osman, K. Chabot, U. T. Sener, J. A. Wilcox, D. Isakov, H. Wang, T. A. Bale, R. Chaligné, J. C. Sun, C. Brown, D. Pe’er, A. Boire, Interferon-γ orchestrates leptomeningeal anti-tumour response. Nature, (2025).

27. P. Bhat, G. Leggatt, N. Waterhouse, I. H. Frazer, Interferon-γ derived from cytotoxic lymphocytes directly enhances their motility and cytotoxicity. Cell Death Dis 8, e2836 (2017).

28. G. S. Reid, X. Shan, C. M. Coughlin, W. Lassoued, B. R. Pawel, L. H. Wexler, C. J. Thiele, M. Tsokos, J. L. Pinkus, G. S. Pinkus, S. A. Grupp, R. H. Vonderheide, Interferon-gamma-dependent infiltration of human T cells into neuroblastoma tumors in vivo. Clin Cancer Res 15, 6602–6608 (2009).

29. S. Zhang, K. Kohli, R. G. Black, L. Yao, S. M. Spadinger, Q. He, V. G. Pillarisetty, L. D. Cranmer, B. A. Van Tine, C. Yee, R. H. Pierce, S. R. Riddell, R. L. Jones, S. M. Pollack, Systemic Interferon-γ Increases MHC Class I Expression and T-cell Infiltration in Cold Tumors: Results of a Phase 0 Clinical Trial. Cancer Immunol Res 7, 1237–1243 (2019).

30. K. Stifter, J. Krieger, L. Ruths, J. Gout, M. Mulaw, A. Lechel, A. Kleger, T. Seufferlein, M. Wagner, R. Schirmbeck, IFN-γ treatment protocol for MHC-I(lo)/PD-L1(+) pancreatic tumor cells selectively restores their TAP-mediated presentation competence and CD8 T-cell priming potential. J Immunother Cancer 8, (2020).

31. C. D. Pham, C. Flores, C. Yang, E. M. Pinheiro, J. H. Yearley, E. J. Sayour, Y. Pei, C. Moore, R. E. McLendon, J. Huang, J. H. Sampson, R. Wechsler-Reya, D. A. Mitchell, Differential Immune Microenvironments and Response to Immune Checkpoint Blockade among Molecular Subtypes of Murine Medulloblastoma. Clin Cancer Res 22, 582–595 (2016).

32. J. Fioravanti, J. Medina-Echeverz, N. Ardaiz, C. Gomar, Z. P. Parra-Guillén, J. Prieto, P. Berraondo, The fusion protein of IFN-α and apolipoprotein A-I crosses the blood-brain barrier by a saturable transport mechanism. J Immunol 188, 3988–3992 (2012).

33. E. F. Spinazzi, M. G. Argenziano, P. S. Upadhyayula, M. A. Banu, J. A. Neira, D. M. O. Higgins, P. B. Wu, B. Pereira, A. Mahajan, N. Humala, O. Al-Dalahmah, W. Zhao, A. V. Save, B. J. A. Gill, D. M. Boyett, T. Marie, J. L. Furnari, T. D. Sudhakar, S. A. Stopka, M. S. Regan, V. Catania, L. Good, S. Zacharoulis, M. Behl, P. Petridis, S. Jambawalikar, A. Mintz, A. Lignelli, N. Y. R. Agar, P. A. Sims, M. R. Welch, A. B. Lassman, F. M. Iwamoto, R. S. D’Amico, J. Grinband, P. Canoll, J. N. Bruce, Chronic convection-enhanced delivery of topotecan for patients with recurrent glioblastoma: a first-in-patient, single-centre, single-arm, phase 1b trial. Lancet Oncol 23, 1409–1418 (2022).

34. J. M. Zaretsky, A. Garcia-Diaz, D. S. Shin, H. Escuin-Ordinas, W. Hugo, S. Hu-Lieskovan, D. Y. Torrejon, G. Abril-Rodriguez, S. Sandoval, L. Barthly, J. Saco, B. Homet Moreno, R. Mezzadra, B. Chmielowski, K. Ruchalski, I. P. Shintaku, P. J. Sanchez, C. Puig-Saus, G. Cherry, E. Seja, X. Kong, J. Pang, B. Berent-Maoz, B. Comin-Anduix, T. G. Graeber, P. C. Tumeh, T. N. Schumacher, R. S. Lo, A. Ribas, Mutations Associated with Acquired Resistance to PD-1 Blockade in Melanoma. N Engl J Med 375, 819–829 (2016).

35. J. Qiu, B. Xu, D. Ye, D. Ren, S. Wang, J. L. Benci, Y. Xu, H. Ishwaran, J. C. Beltra, E. J. Wherry, J. Shi, A. J. Minn, Cancer cells resistant to immune checkpoint blockade acquire interferon-associated epigenetic memory to sustain T cell dysfunction. Nat Cancer 4, 43–61 (2023).

36. J. Eng, E. Bucher, Z. Hu, M. Sanders, B. Chakravarthy, P. Gonzalez, J. A. Pietenpol, S. L. Gibbs, R. C. Sears, K. Chin, Robust biomarker discovery through multiplatform multiplex image analysis of breast cancer clinical cohorts. bioRxiv, (2023).

37. J. Eng, E. Bucher, Z. Hu, T. Zheng, S. L. Gibbs, K. Chin, J. W. Gray, A framework for multiplex imaging optimization and reproducible analysis. Commun Biol 5, 438 (2022).

