## Supplementary Material for "Interferon Restores Antigen Presentation and Sensitizes Medulloblastoma to T Cell Killing"

### List of Supplementary Materials

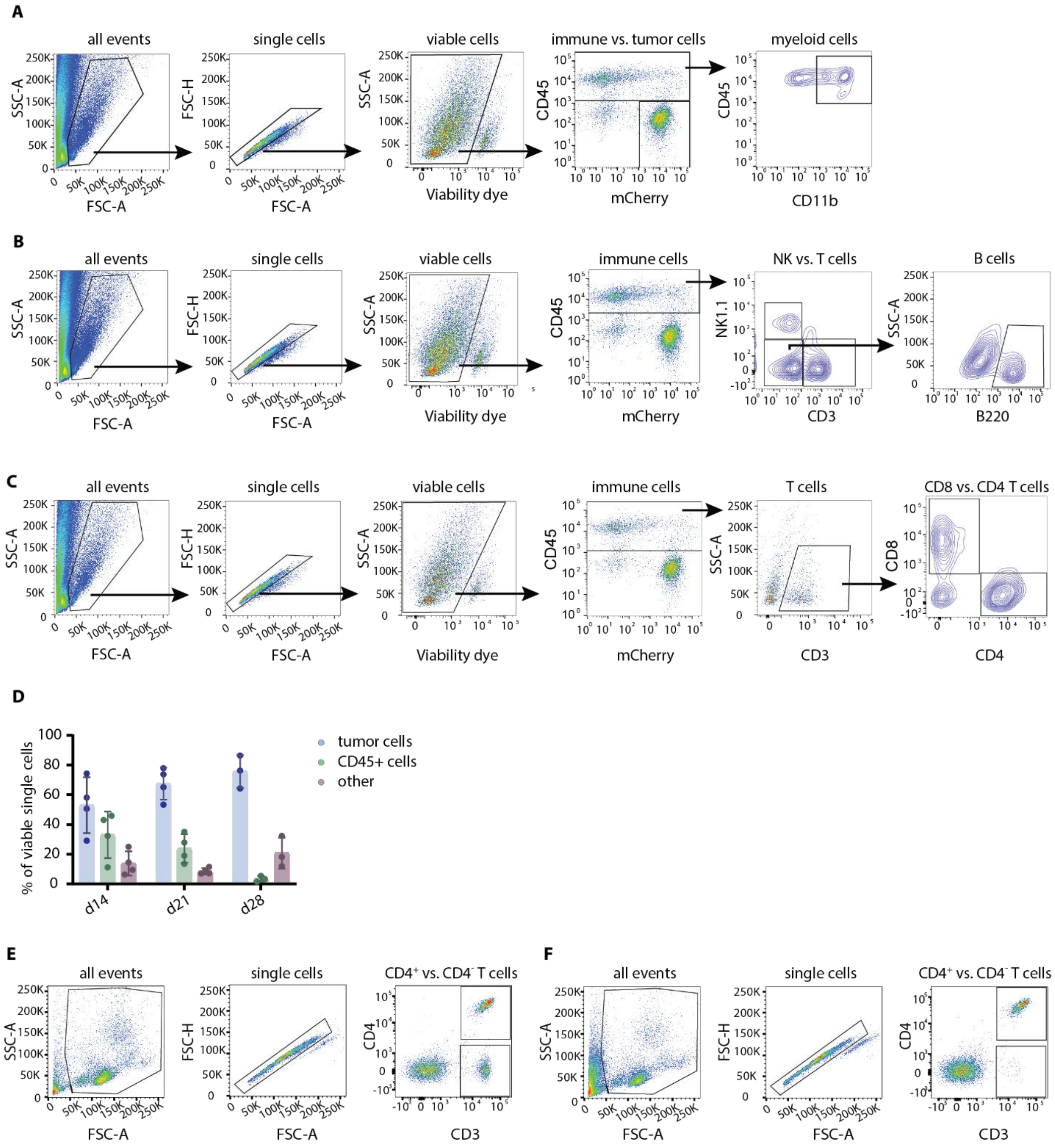

**Fig. S1 Tumor infiltrating immune cells.**

Gating strategies for quantification of (A) myeloid cells and mCherry positive tumor cells, (B) NK cells, T cells and B cells, and (C) CD4 and CD8 T cells. (D) Quantification of tumor cells, CD45 positive immune cells and other cells at 14, 21, and 28 after tumor cell transplantation. (E, F) Gating strategy for analysis of CD8 T cell depletion. As the depletion antibody may impede the binding of fluorescently labeled anti-CD8 antibody, CD8 T cells were assessed in (E) control animals and (F) treated animals by expression of CD3 and lack of CD4.

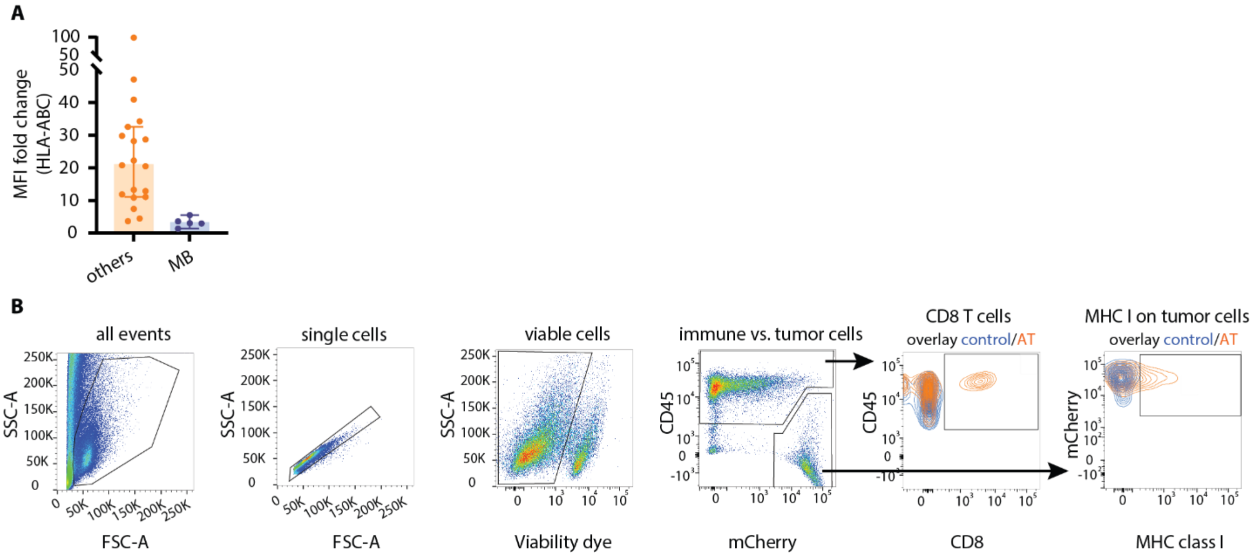


**Fig. S2 Expression of MHC class I by medulloblastoma cells.**

(A) Expression level of HLA-ABC of pediatric brain tumors (n = 20) and medulloblastoma (n = 5) indicated by the fold change of the mean fluorescence intensity compared to isotype antibody control. (B) Gating strategy to detect CD8 T cells transferred to tumor-bearing recipient mice, and the expression of MHC class I on tumor cells.

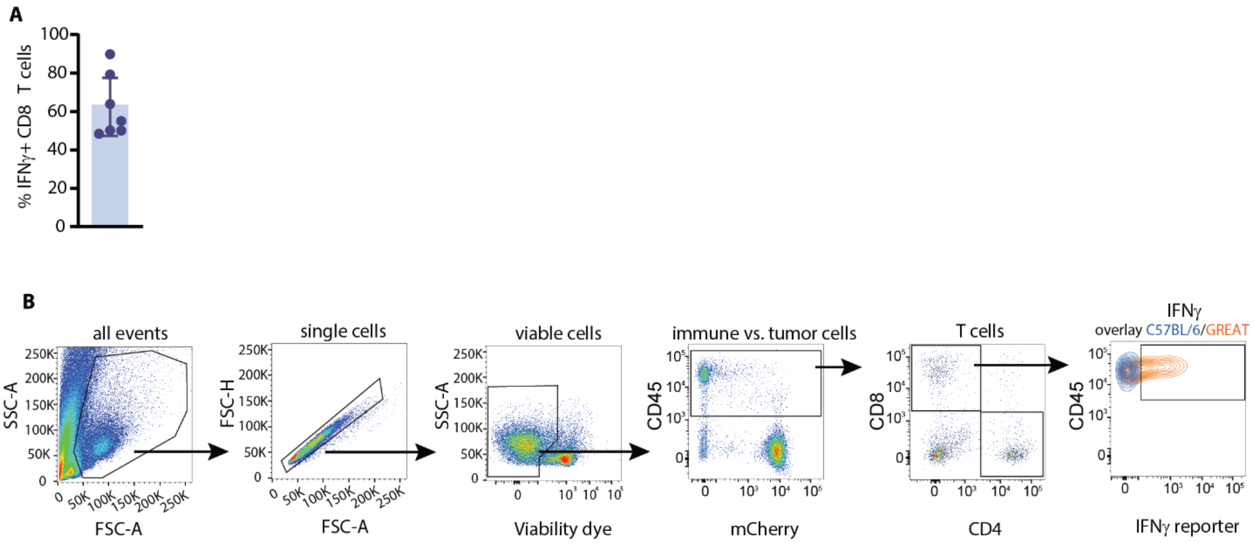


**Fig. S3 Tumor infiltrating CD8$^{+}$ T cells express IFNγ**

(A) Quantification of tumor infiltrating IFNγ positive CD8$^{+}$ T cells, 14 days post tumor cell transplantation. GREAT mice were used as hosts which have an IFNγ promoter that drives an IRES-eYFP reporter expression to enable analysis of T cell activation, n = 7; mean and 95% CI. (B) Gating strategy for quantification of IFNγ expression by CD8+ T cells.

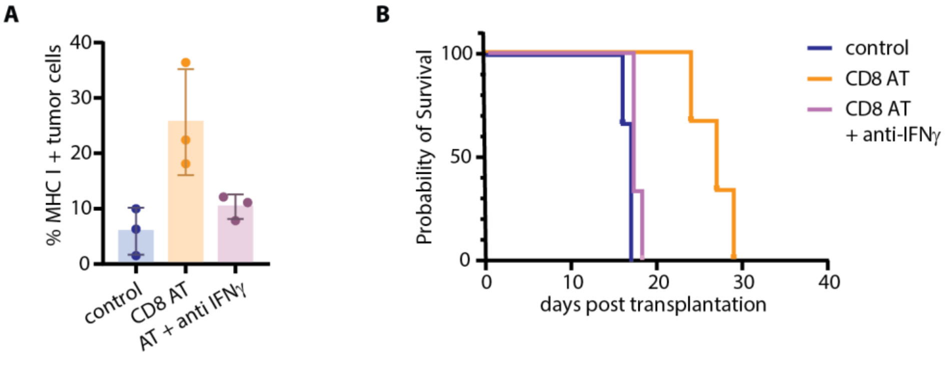


**Fig. S4 IFNγ blockade impedes MHC class I upregulation**

(A) Percent MHC class I positive (OVA positive) tumor cells isolated from NSG control mice, mice that received OT-I CD8$^{+}$ T cells (AT), or mice that received IFNγ blocking antibody in addition to CD8$^{+}$ T cells, analyzed by flow cytometry (n = 3). (B) Corresponding survival curve, n = 3.

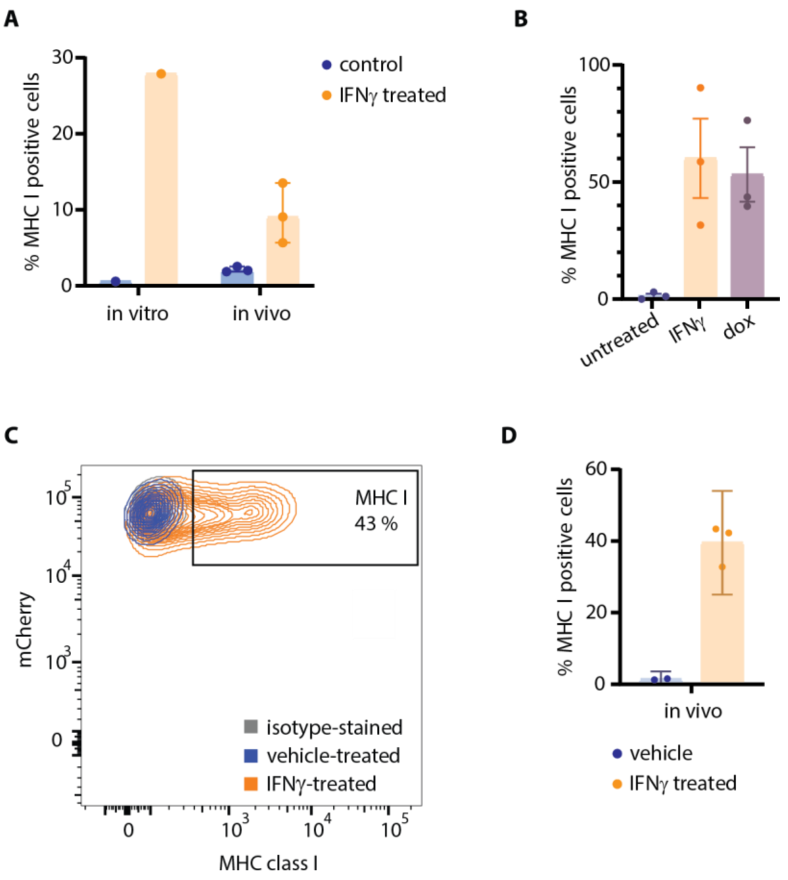


**Fig. S5 IFNγ treatment in vivo and in vitro.**
(A) Flow cytometric quantification of tumor cells treated in vitro, and isolated from control and systemically IFNγ treated NSG mice. Animals were injected i.p. with 40µg daily for 2 consecutive days. (B) In vitro treatment of MP-TetOne cells with 10ng/ml IFNγ or 2mg/ml doxycycline for 72h followed by flow cytometric analysis of MHC class I expression. (C) Flow cytometry contour plot of tumor cells stained for MHC class I or isotype control and (D) corresponding quantification. Cells were freshly isolated from NSG mice treated for 3 days with vehicle (n = 2) or IFNγ (n = 3) via convection-enhanced delivery (CED).
